# Spontaneous emergence of multicellular heritability

**DOI:** 10.1101/2021.07.19.452990

**Authors:** Seyed Alireza Zamani Dahaj, Anthony Burnetti, Thomas C. Day, Peter J. Yunker, William C. Ratcliff, Matthew D. Herron

**Author notes:** Corresponding authors; William C. Ratcliff’s; Matthew D. Herron’s.

## Abstract

The Major Transitions in evolution include events and processes that result in the emergence of new levels of biological individuality. For collectives to undergo Darwinian evolution, their traits must be heritable, but the emergence of higher-level heritability is poorly understood and has long been considered a stumbling block for nascent evolutionary transitions. A change in the means by which genetic information is utilized and transmitted has been presumed necessary. Using analytical models, synthetic biology, and biologically-informed simulations, we explored the emergence of trait heritability during the evolution of multicellularity. Contrary to existing theory, we show that no additional layer of genetic regulation is necessary for traits of nascent multicellular organisms to become heritable; rather, heritability and the capacity to respond to natural selection on multicellular-level traits can arise “for free.” In fact, we find that a key emergent multicellular trait, organism size at reproduction, is usually more heritable than the underlying cell-level trait upon which it is based, given reasonable assumptions.

## Introduction

Life on Earth is, and always has been, predominantly unicellular. At various times, unicellular populations have evolved multicellular structures, bodies, or thalli. In some cases, these now multicellular populations massively diversified, giving rise to the major radiations of multicellular life: plants, animals, fungi, and various groups of seaweeds. These radiations have massively modified the biosphere, ultimately affecting the ecology and evolution of multicellular and unicellular organisms alike.

Our focus here is on the first steps in the transition to multicellular life: the evolution of multicellularity *per se* and the early evolution of nascent multicellular populations. A necessary early step in the transition from unicellular to multicellular life is the evolution of a mechanism that keeps cells together (Abedin and King 2010; Niklas and Newman 2013; Tarnita et al. 2013). In some cases, the population may have already expressed a multicellular phenotype plastically, as is known to happen in some algae (Iwasa and Murakami 1968; Kapsetaki et al. 2016) and choanoflagellates (Alegado et al. 2012), and the transition in question is from facultative to obligate multicellularity. In other cases, the multicellular phenotype may be entirely novel, resulting in a transition from a purely unicellular population to an obligately multicellular one. Both processes have been observed in microbial evolution experiments in which a selective pressure was applied to unicellular laboratory populations (Becks et al. 2010; Herron et al. 2019; Ratcliff et al. 2012, 2013).

The resulting populations consist of clusters of clonally related cells. These clusters have traits that did not exist in the unicellular population, for example cluster diameter, cluster shape, and number of cells per cluster. For selection to act on emergent multicellular traits, however, they must be heritable.

The origin of multicellular heritability has widely been viewed as a problem for evolutionary transitions to multicellular life (Libby and Rainey 2013; Michod 2000; Michod and Roze 2001; Roze and Michod 2001; Simpson 2011; Watson et al. 2016). Specifically, prior work has argued that nascent multicellular organisms must ‘reorganize’ the relationship between genotype and multicellular phenotype (i.e., through the evolution of a developmental program), providing a heritable basis for multicellular traits (Maynard Smith and Szathmáry 1995; Michod 2000). Here, we consider the heritability of multicellular traits in a laboratory-derived multicellular organism that has not yet had time to evolve multicellular adaptations.

Our approach relies on the fact that all cluster-level traits can, in principle, be expressed as functions of cell-level traits, though in many cases these functions will be so complex as to be intractable (Herron et al. 2018). They may, for example, include cell-cell communication, complex feedbacks, and interactions with the environment, such that cell-level phenotype is only one of several arguments. For simplicity, we focus on an emergent cluster-level trait whose value is a function of a single cell-level trait: cluster size at division.

In multicellular ‘snowflake’ yeast, the size that clusters can grow to before reproducing (see Supplementary Movie 1) depends on the biophysics of cellular packing. The evolution of more elongate cells (i.e., cells with a higher aspect ratio) reduces the density at which cells pack in the group, thus increasing the size to which clusters grow before physical strain drives cluster reproduction (Jacobeen et al. 2018a,b). Through a combination of analytical modeling, experimental manipulation and simulations, we show that the heritability of cluster size at reproduction exceeds that of the underlying cell-level trait, cellular aspect ratio, across a wide range of conditions. Rather than requiring a ‘reorganization’ of multicellular heredity (Michod 2000), these results demonstrate how the first step in the transition to multicellularity, group formation, can itself generate novel multicellular traits that are remarkably heritable.

## Results

### Analytical model

The breeder’s equation of quantitative genetics shows that a trait’s response to selection (*R*) is the product of the selective coefficient (*S*) and the heritability of the trait under selection (*h*^2^): *R* = *h*^2^*S*. Heritability in this formulation is the narrow-sense heritability, the proportion of phenotypic variation explained by additive genetic variation, and this formulation predicts the response to a given strength of selection in an obligately sexual population (Lynch et al. 1998). For simplicity, we consider asexual populations, for which the relevant measure of heritability is broad-sense heritability, *H*^2^, the proportion of phenotypic variation explained by all genetic variation (Lynch et al. 1998). The response to selection in an asexual population, then, is *R* = *H*^2^*S* (Lynch et al. 1998).

Thus we could, in principle, predict the response of a nascent multicellular population to a given strength of selection if we could predict *H*^2^. Doing so is not trivial, though. Heritability is not an inherent property of organisms or species. Rather, it is a population-level statistical measure that applies only to a particular trait in a particular population at a particular time.

Prior biophysical modeling (which we confirm experimentally, see Figure 2) suggests there is a linear relationship between the size to which clusters grow before they reproduce and cellular aspect ratio (Jacobeen et al. 2018a). Snowflake yeast grow as branched, tree-like groups of cells. When their cells are more elongate, this reduces cellular crowding within the interior of the cluster, slowing the rate of stress accumulation that ultimately causes the cluster to fracture. As a result, genotypes with a higher cellular aspect ratio are capable of forming larger clusters. Using this relationship as a starting point, we developed a theoretical model for describing the relationship between the heritability of cellular aspect ratio, a trait that is directly affected by mutations in cell cycle genes, and the number of cells in the cluster at division, which is an emergent property of cellular aspect ratio.

**Figure 1:**
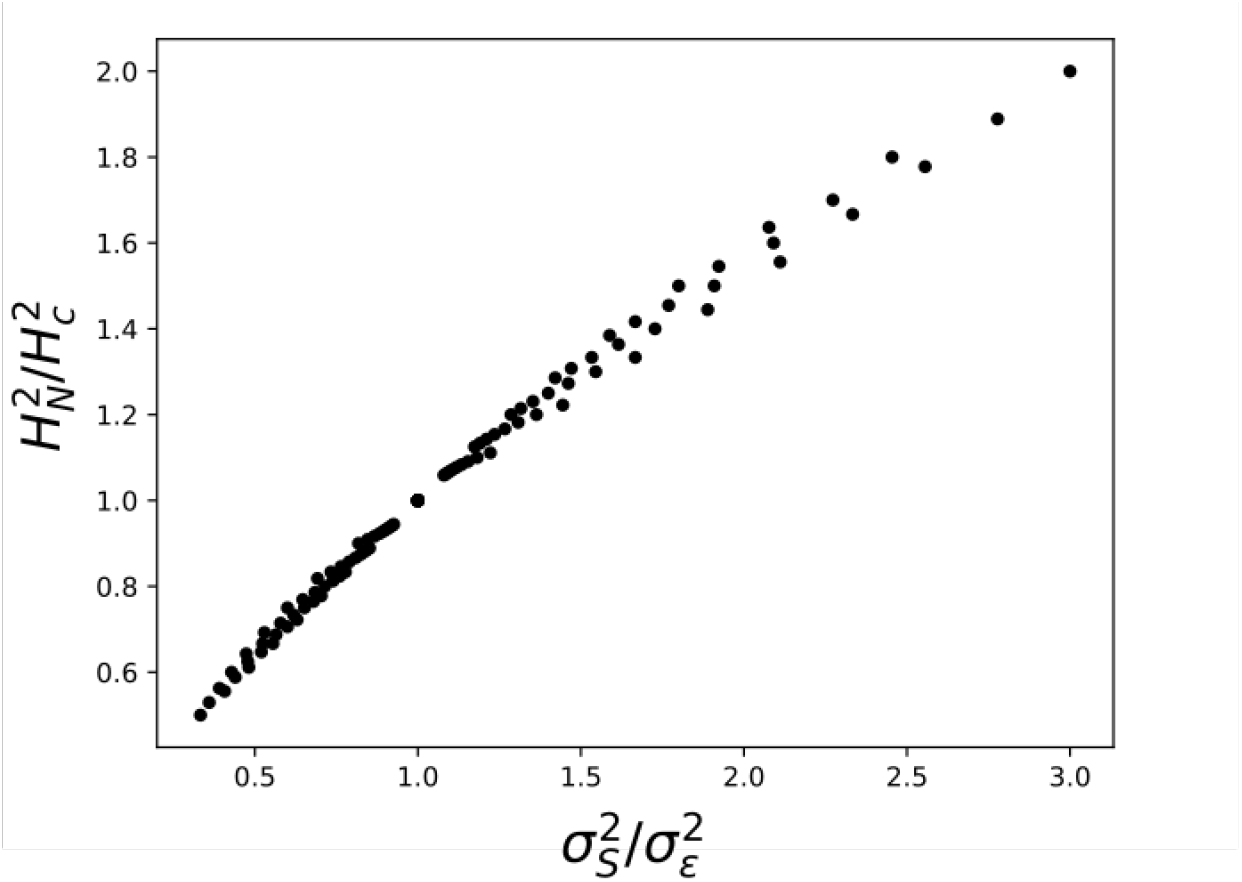
The heritability of a group-level trait, number of cells at division 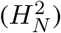, will be higher than that of a cell-level trait, aspect ratio 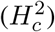, whenever cellular developmental noise 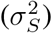 is higher than collective level developmental noise 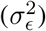.

**Figure 2:**
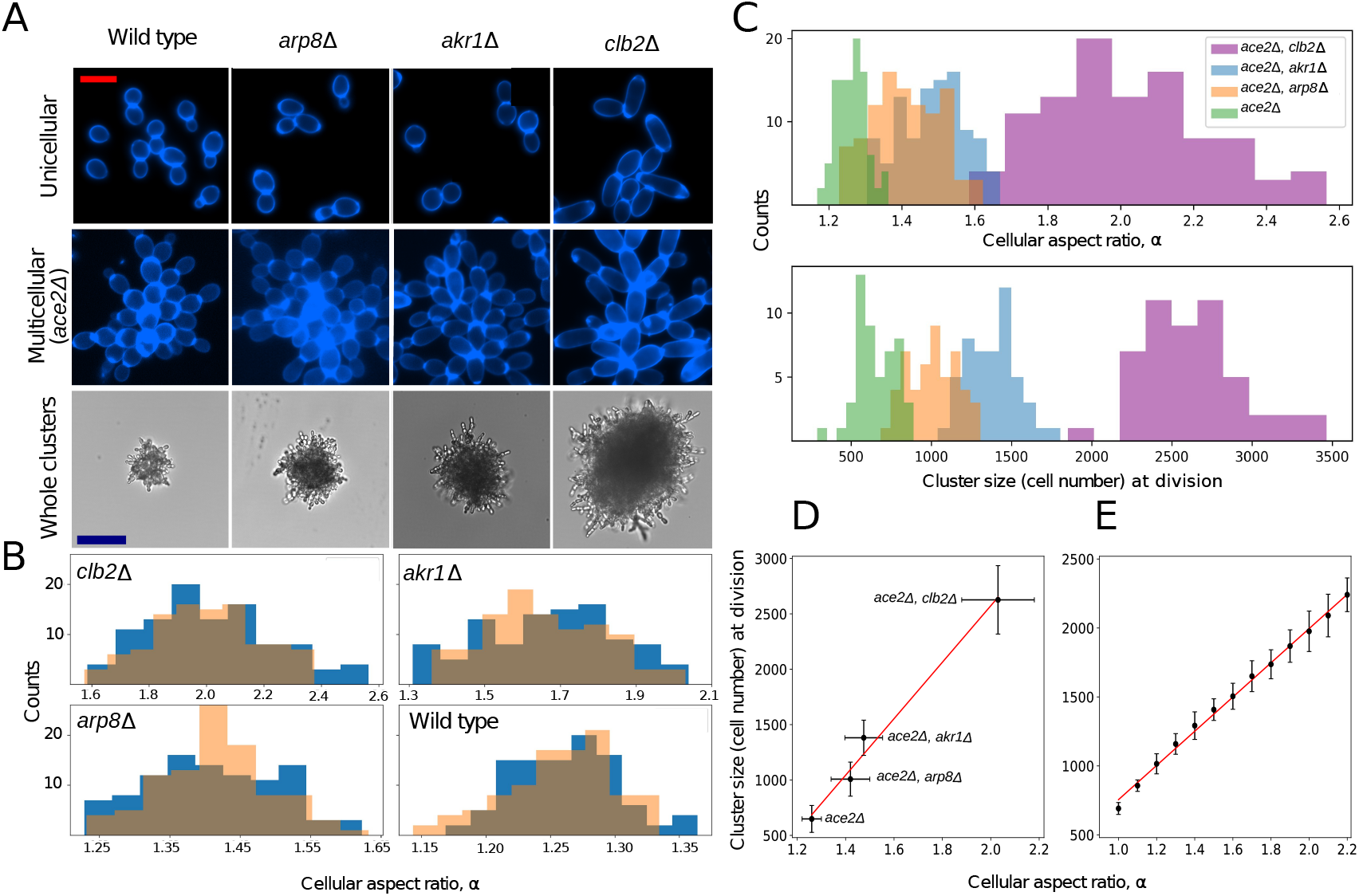
Experimentally modifying cellular aspect ratio. A) Cell cycle mutants with functional (top row) or nonfunctional (second row) *ACE2* alleles. The bottom row shows representative brightfield images of multicellular *ace2*∆, demonstrating how cellular elongation increases cluster size. Scale bar in the top two rows of A is 10 *μm*, and in the bottom row of A is 50 *μm*. B) Distribution of cell aspect ratio for each genotype with functional *ACE2* (orange bars) and nonfunctional *ace2* (blue bars). C) Distribution of cellular aspect ratio (top) and number of cells at fracture (bottom) for the four multicellular genotypes in our experiment. Cluster size at division is an approximately linear function of cellular aspect ratio. Empirical data in (D) (*y* = 2549.8*x* − 2526.6, *r*^2^ = 0.96) and results from a biophysical simulation of cluster growth and fracture in (E) (*y* = 1240*x* − 485, *r*^2^ = 0.99).

First, we calculated broad-sense heritability as the correlation coefficient between parental phenotype *α* (cellular aspect ratio) and offspring phenotype *α*′ for cellular aspect ratio, partitioning the phenotype into its key components (Fisher 1919):

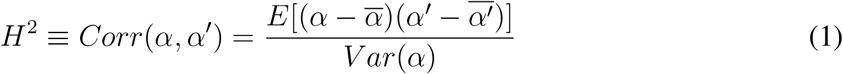

Where:

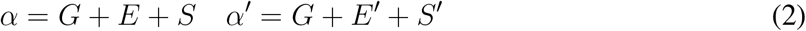

In this equation *G* is the genetic mean of aspect ratio, *E* is environmental effects, and *S* is intrinsic variation in the expression of the trait, which we refer to as developmental noise. Here we assume that *S* is a white noise, with mean zero and variance 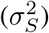.

If there is no temporal correlation except for the same genotype between offspring and par ts, then 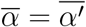 and we can rewrite the numerator as:

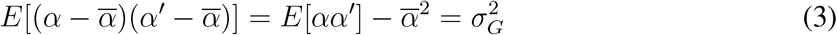

So the heritability of aspect ratio is equal to:

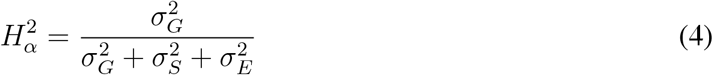

If *N* is a linear function of the mean of cellular aspect ratio, then *N* is also a linear function of *G*, because all the cells in a clonal cluster have the same genetic mean value for aspect ratio. But because developmental noise *S* is random and different for each cell, its expected value for all the cells in a cluster is zero. We also consider developmental noise at the cluster level, *ϵ*, which accounts for internally-generated (i.e., not environmental) variation in cluster phenotype. So for *N* we have: *N* = *a*(*G* + *E* + *ϵ*), 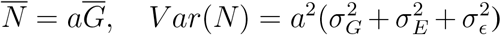. Given these equations, the heritability of the number of cells at division in the simplest form is equal to:

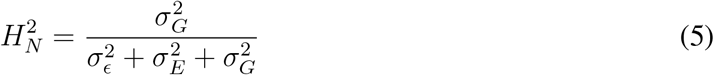

so the ratio of the heritability of the number of cells per cluster and cell size is equal to:

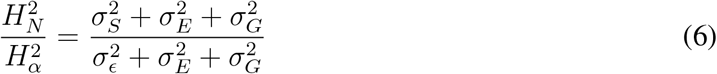

Thus, in principle, an emergent multicellular trait (cluster size at reproduction) can not only be heritable prior to the evolution of multicellular developmental regulation, but can even be more heritable than the the underlying cellular trait as long as 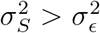 (Figure 1).

### Experimental system

We experimentally examined the relative heritability of mutations affecting both a cellular trait (cellular aspect ratio) and an emergent multicellular trait (number of cells in the cluster at reproduction) using the snowflake yeast model system (Ratcliff et al. 2012). We deleted three cell cycle regulatory genes (*AKR1, ARP8, CLB2*), generating a range of mutants that varied in their mean cellular aspect ratio. In each case, we generated otherwise isogenic unicellular and multicellular versions of these strains by leaving *ACE2* functional, or deleting the *ACE2* open reading frame, respectively (Ratcliff et al. 2015) (Figure 2A).

In both unicellular and multicellular isolates, we measured the cellular aspect ratio (ratio of length to width) of cells that were a single division old. Because cellular aspect ratio varies with replicative age in yeast (Jin et al. 2019), this provides an upper bound estimate of the heritability of this trait at the cell-level by excluding age-dependent phenotypic variation. *arp8*∆, *akr1*∆, and *clb2*∆ increased aspect ratio from a mean of 1.26 in the ancestor, to an average of 1.41, 1.49 and 2.02, respectively (Figure 2B; *F_3,399_* = 642.7, *p* < 0.0001, ANOVA, all means significantly different at *p* = 0.001 with Tukey’s HSD test). Whether or not Ace2p was active did not significantly affect cellular aspect ratio in any of our genotypes (Figure 1B, *t* = −0.96, *p* = 0.33 for the wild type, *t* = 1.12, *p* = 0.26 for *arp8*∆, *t* = −.69, *p* = 0.49 for *akr1*∆, *t* = −0.29, *p* = 0.76 for *clb2*∆; two-tailed t-tests).

To measure the number of cells in the cluster at reproduction (*N*) we first measured the packing fraction (*ϕ*) of each genotype and its size just before division (*V_c_l*) using time-lapse microscopy. We also measured the volume of cells (*v_c_e*) for each genotype, and using the formula *ϕ* = *Nv_c_e/V_c_l*, calculated the number of cells in the cluster at the point of fracture (Figure 2C). This is a key life history trait of snowflake yeast, determining their maximum size. Cluster size at reproduction is an approximately linear function of cellular aspect ratio (*y* = 2549.8*x* − 2526.6, *r*^2^ = 0.96, Figure 2D). To more comprehensively examine the functional form of this relationship (our experiments are limited to four genotypes), we adapted the 3D biophysical simulation from (Jacobeen et al., 2018a), which allows us to simulate snowflake yeast growth until packing strain arising from cellular division causes fracture (Figure 2E). Simulating the growth and fracture of 25 clusters for cellular aspect ratios ranging from 1-2.2 (in increments of 0.1), we confirm that group size at reproduction indeed is an approximately linear function of cellular aspect ratio (*y* = 1240*x* − 485, *r*^2^ = 0.99).

### Quantifying heritability

A trait’s heritability is a statistical feature of a population, not an inherent property of an organism, and is highly sensitive to population composition (Fisher 1919). We used a variance partitioning approach (Lynch et al. 1998) to calculate the broad-sense heritability of cellular aspect ratio and group size at division from our experimental data (see the Supplementary Material for a formal derivation). Because each cell cycle mutant has a different genetic mean cellular aspect ratio, the heritability of both the cell and group-level traits will depend on the frequency of each genotype in the population (equation 4). Therefore, we measured the heritability of both cellular aspect ratio and cluster size at reproduction by simulating all possible hypothetical populations, generating populations of 1000 individuals by bootstrapping, without replacement, from our experimental data (Figure 3). Because it is difficult to visualize four-dimensional space, which is requried if we simultaneously vary the frequency of four genotypes, we examined all possible three-member populations, varying the frequency of each genotype from 2.5 − 95%. Setting a maximum genotype frequency of 95% ensures that the population never becomes monomorphic, a pathological case where heritability is undefined. In nearly all possible populations, the multicellular trait, number of cells in the group at reproduction, is more heritable than the underlying cell-level trait, cellular aspect ratio, despite the fact that cellular aspect ratio is the only trait being modified directly via mutation (Figure 3).

**Figure 3:**
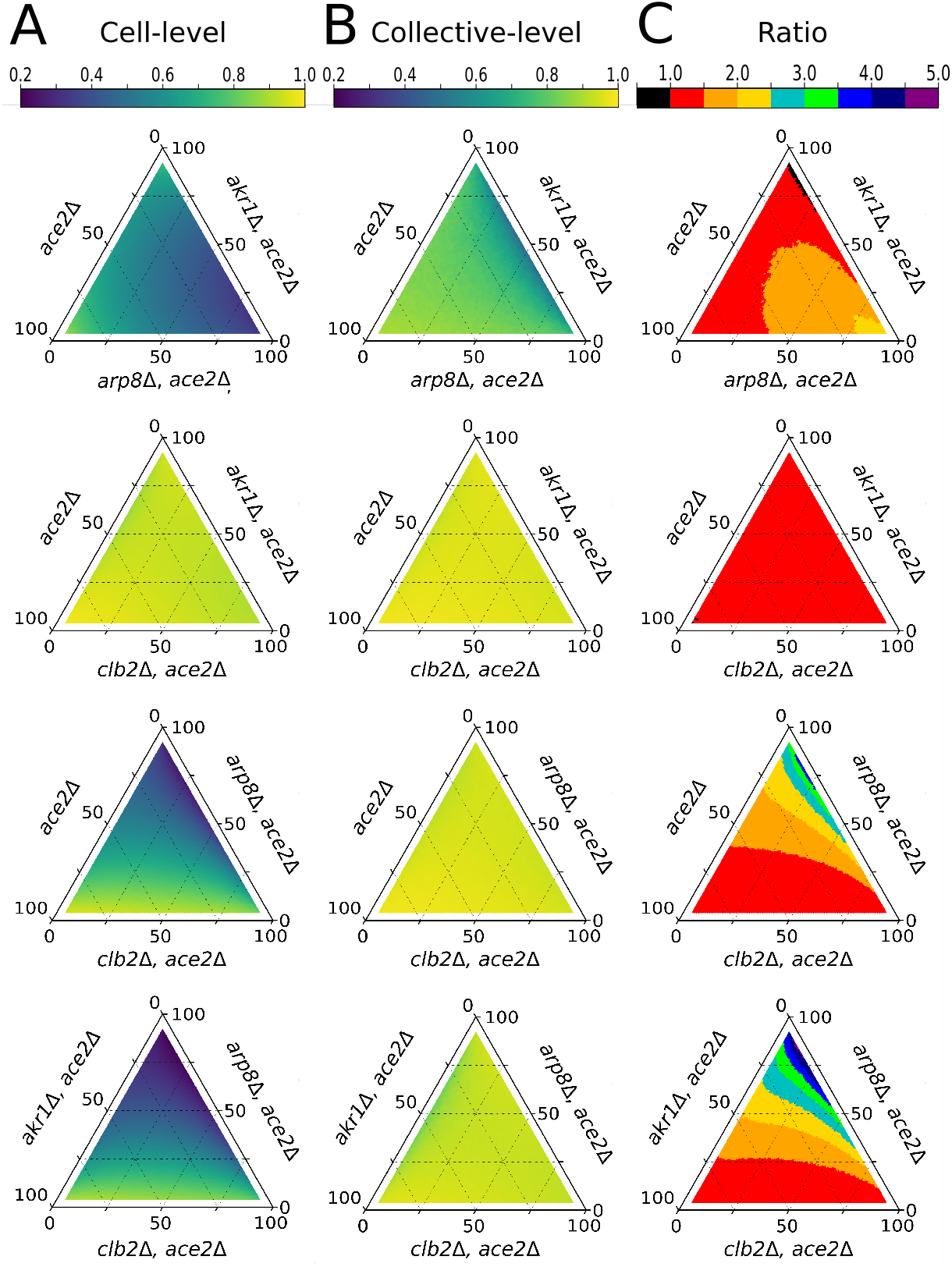
The heritability of an emergent multicellular trait exceeds that of the corresponding cell-level trait over a wide range of simulated populations. Each triangle plot represents the heritability of the cell-level trait, cellular aspect ratio (A), collective-level trait, cluster size at reproduction (B), or the ratio of collective to cell-level trait heritabilities (C). Shown are all possible 3-way combinations of the four genotypes (each row being a different population composition), across all possible population frequencies ranging from 2.5-95% per genotype.

## Discussion

During a major evolutionary transition, organisms form novel collectives that become more complex and biologically individuated by gaining adaptations (Herron 2021). For adaptations to occur at the collective level, nascent collectives must not only be capable of reproduction, but must also possess heritable variation in traits that affect fitness. In this paper, we focus on the origin of heritability during the transition to multicellular life, showing that the evolution of developmental systems that ‘reorganize’ the relationship between genotype and multicellular phenotype (Maynard Smith and Szathmáry 1995; Szathmáry and Maynard Smith 1995) are not required for its emergence. Instead, we confirm predictions by a previous analytical model (Herron et al. 2018), that the heritability of the multicellular trait can in fact be higher than that of the underlying cellular trait, even in the absence of subsequently-evolved developmental regulation. Although the ratio of cellular and multicellular heritabilities depends on the proportion of each genotype in the population, it is in nearly all cases greater than one.

While it might seem surprising that an emergent multicellular trait can be more heritable than a cell-level trait directly encoded for by genes, multicellular groups possess a powerful advantage over isolated cells: the ability to average out stochastic variation in cellular phenotype (Herron et al. 2018). Because of this averaging effect, cluster phenotypes are less variable within a genotype than cell phenotypes, so non-genetic variation is a smaller proportion of total phenotypic variation (Herron et al. 2018).

Although it is routinely estimated and widely recognized as useful in predicting responses to selection in clonally reproducing multicellular organisms (Bokmeyer et al. 2009; Bonos et al. 2003; Burton and Devane 1953; De Meester 1989; Oard et al. 1983; Stratton 1991; Vorburger 2005; Weisser and Braendle 2001; Yampolsky and Ebert 1994), *H*^2^ is typically assumed to be 1 in asexually reproducing microbes (Baselga-Cervera et al. 2016; Bouma and Lenski 1988; Dykhuizen 1990; Gerstein et al. 2011; Gullberg et al. 2011; López Rodas et al. 2006). This assumption is so widespread that the concept of heritability is rarely addressed in discussions of microbial evolution; rather, the response to selection is simply assumed equal to the strength of selection (*R* = *S*). However, *H*^2^ is never 1 for a continuously varying trait in a real population. Because broad-sense heritability is defined as the ratio of genetic variance to total phenotypic variance, which includes genetic variance, the only way for *H*^2^ to equal 1 is for there to be zero non-genetic variance.

In reality, phenotypic heterogeneity is widespread even among genetically identical individuals reared in carefully controlled environments. This phenomenon is well documented among microbes, but it is also true for diverse plants, fungi and animals (Vogt 2015) and for traits as diverse as gene expression (Ballouz et al. 2019; Elowitz et al. 2002), antibiotic (Balaban et al. 2004) and formaldehyde (Lee et al. 2019) resistance, motility (Salek et al. 2019), phototaxis (Kain et al. 2012), cell number (Herron et al. 2014), predator evasion behavior (Schuett et al. 2011), and number of sense organs (Vogt et al. 2008).

While the emergence of multicellular heritability is widely regarded as a significant challenge in this major evolutionary transition (Doulcier et al. 2020; Okasha 2003, 2013), we show that multicellular heritability may in fact arise spontaneously, as a side-effect of group formation. Our experimental design excludes the possibility of a subsequently-evolved layer of regulation, since the unicellular strains are genetically identical to the multicellular strains, aside from the *ace2* deletion.

If it were true that heritability of multicellular traits only arose as the outcome of some adaptive process subsequent to the origin of multicellularity, this would, as previous authors have pointed out (Maynard Smith and Szathmáry 1995; Michod 2000; Szathmáry and Maynard Smith 1995), engender a chicken-and-egg problem for the evolution of complex multicellularity: how could heritability arise as an adaptation when heritability is a prerequisite for adaptation? We have shown analytically and empirically that no such problem exists. Multicellular traits are heritable as soon as they come into existence, and this is a straightforward result of the way variation is partitioned. Adaptive modification of multicellular-level traits is thus bootstrapped into feasibility, and subsequent adaptations can build upon earlier ones. We speculate, and indeed see no reason to doubt, that the spectacular complexity that characterizes a few multicellular clades arose through a sequence of adaptive changes that started with a process similar to the one we have described.

## Methods

### Strain construction

All yeast strains were produced from a Y55 strain background made homozygous at all loci via sporulation and selfing, as previously used by the Ratcliff laboratory (Ratcliff et al. 2015). All strains were based on the GOB8 strain bearing a homozygous deletion of the *ACE2* open reading frame with the KanMX cassette, resulting in an *ace2*∆:KanMX / *ace2*∆:KanMX, producing small constitutively multicellular snowflake yeast. Cell-lengthening mutations were identified from previous work showing that disruption of *CLB2* results in an alteration of filamentous growth (Edgington et al. 1999), and screens of genes known to alter the aspect ratio or filamentous growth form of yeast cells (Watanabe et al. 2009).

Strains (Table S1 were produced via standard yeast genetics protocols. Cells were transformed via the lithium acetate + polyethylene glycol transformation method (Gietz and Woods 2006). All deleted genes were removed via transformation using PCR products of plasmid pYM25 (Janke et al. 2004), creating ends-out gene deletion cassettes bearing the hygromycin-resistance hphNTI gene with 50 base pairs of flanking sequence just outside of each reading frame. Sequences can be seen in Table S2. Upon selection of heterozygous deletion strains on Yeast Peptone Dextrose (YPD) + hygromycin plates, colonies heterozygous for the deletion were sporulated and tetrads dissected onto YPD + hygromycin plates, and spores allowed to self-mate before single-colony purification for analysis.

### Cell culture

Yeast were cultured in Yeast Extract Peptone Dextrose (YEPD) media (1% yeast extract, 2% peptone, 2% dextrose) at 30°C. In large-diameter tubes (25mm), each strain was cultured 10 mL YEPD with rapid mixing (250 rpm).

### Measuring cellular packing fraction

To measure the packing fraction of each genotype, we took 1 mL of each cell culture and stained them with DAPI using the following protocol: first we transferred 500 *μL* of each cell culture to a 1.5 ml microcentrifuge tube, and replaced the YEPD media with 500 *μL* of 70% ethanol. Then we shook the tubes at 1300 rpm for 5 minutes at 25°C and removed the ethanol by washing, pelleting and resuspending in 1% phosphate buffered saline solution. We added 0.1% DAPI stock solution (1.25 *mg/*500*ml*) and vortexed vigorously. We incubated the cells for 5 minutes at 25°C, then diluted the stained cells 1:10, and imaged individual clusters without lateral compression on a Nikon Eclipse Ti under brightfield illumination using a 10x objective. From this we calculated the effective radius of each cluster as:

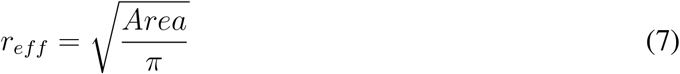

We approximated the volume of each cluster by calculating the volume of a sphere with *R* = *r_eff_*. We then placed a coverslip over the cluster and applied pressure until it was a cellular monolayer, allowing us to image DAPI-stained nuclei using a 20x objective. This allowed us to count the number of cells per cluster (*N_cell_*) using ImageJ-FIJI. We approximated the volume of 100 cells (*V_cell_*) for each genotype (ensuring each was just once cellular generation old) by calculating the volume of a prolate ellipsoid that its minor and major axis are equal to short and long axis of a cell respectively. Using these information and the formula:

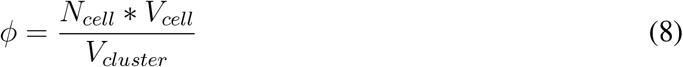

### Quantifying cellular aspect ratio

To measure cellular aspect ratio, we grew samples both with and without functional *ACE2* alleles (i.e., uni- and multicellular versions of these genotypes) overnight at 30°C in YEPD, shaking at 250 RPM. After that, we grew these samples for 2 days until they reached stationary phase. Next, we used calcofluor-white to stain cell walls, incubating them in the dark for 30 mins in a one micromolar solution, and imaged them with a Nikon Eclipse Ti microscope under ultraviolet excitation for blue fluorescence using a 20x objective. We measured the aspect ratio of 50 individual cells for the ancestor and each cell cycle mutant, repeating the experiment both with and without functional *ACE2* strains. Cell boundaries were identified manually using ImageJ-FIJI. In all cases we only considered cells with a single bud scar, mitigating issues stemming from possible age-dependent variation in cellular aspect ratio.

### Quantifying cluster size at division

We determined the size of clusters at division using timelapse brightfield microscopy. Clusters were inoculated into 1*μl* drops of YEPD medium and grown overnight, and imaged every five minutes at using a 4x objective. Using ImageJ-FIJI, we then measured the radius of 50 clusters per each genotype in the timestep just before multicellular division. Using the formula described above, we calculated the volume of each cluster, and, knowing mean cell size and packing fraction of these genotypes, were able to estimate the number of cells in each group just prior to division.

### Biophysical simulations

We simulated snowflake yeast by adapting the spatially explicit approach described in (Jacobeen et al. 2018a,b). In these simulations, cells were modeled as prolate ellipsoids. Briefly, new cells were procedurally added to the surface of existing cells in the cluster, resulting in cluster growth in a manner analogous to the way snowflake yeast grow. Each cell was allowed to divide as long as there was space for an offspring cell. All elements of the simulation were held constant for Figure 2D, varying only the input distributions of cellular aspect ratio. See the following section for a more detailed description of the simulation.

#### Generating snowflakes

New cells are added in an analogous manner to cellular growth in actual snowflake yeast. Every generation, each cell in the cluster attempts to divide. They first bud from their distal pole (i.e. *θ* = 0 ± 10 degrees), with subsequent cells budding at a polar angle *θ* = 45 degree, and randomly chose an azimuthal angle from a uniform distribution *ϕ* ∈ [0, 2*π*]; in other words, after the first bud, cells generally appeared along a “budding ring” at the distal pole of the cell. There was a 20% chance that the first bud would appear along this budding ring instead of exactly at the pole. After 3 bud scars, there was a 50% chance that new cells bud on the proximal side (*π* − *θ*) instead of the distal side. If new bud scar locations were located within 1.17 microns of a previous bud scar on the cell surface, then the new bud was unable to form and the mother cell lost its chance to generate a new cell for that generation. The orientation of the new cell is determined by the surface normal to the mother cell at the position of the bud site. The cellular aspect ratio of each new bud was drawn from a normal distribution centered on the mean value *α* with standard deviation *α*/10.

After each generation of cell division, there is a chance that two cells overlap due to the proximity of their bud scars, or because two cells grew into the same space, etc. The amount by which any two cells overlap was computed by *d* = *D* − *R* if *d* < 0, where *d* is the overlapping distance, *D* is the distance between the centers of two cells, and *R* = *R*_1_ + *R*_2_ is the sum of the two cell’s equatorial radii. We modeled the energy associated with this overlap as Hertzian (i.e. *u* ∼ *d*^5*/*2^). The total energy of the overlaps is then *U* = Σ_*i*_ *u* where *i* runs over all pairs of cells. The total overlapping energy was recorded after each generation of cell division, and if there were more than 20 budding events, after every 20 budding events. Once the total overlapping energy succeeded a threshold amount *U_overlap_*, the simulation was stopped. This was the final size of the cluster. 25 groups were simulated in this manner for each aspect ratio.

## Acknowledgements

This work was supported by grants from NSF (DEB-1723293 and DEB-1845363), NIH (GM138030) and NASA (NNA17BB05A). This material is based upon work while M.D.H. was serving at the National Science Foundation.

## Correspondence

Correspondence and requests for materials should be addressed to W.C.R. and M.D.H.

## Supplementary material

### Calculating heritability by partitioning variance. Clonal population

Let’s assume that we have *N* clonal lineages and each has *n_i_* members where *i* is from 1 and *N*. For the simplest case we can assume a linear model for the phenotype of each individual:

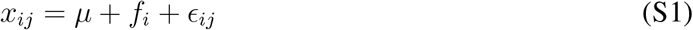

*x_ij_*: The phenotype of the *j*th member of the *i*th family.

*f_i_*: The effect of the *i*th family where: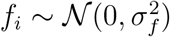

*ϵ_ij_*: The residual and random effect dues to all other variables: 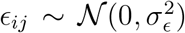 From which we have:

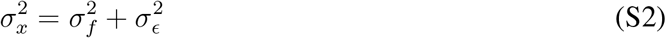

We can partition the total variation on phenotype in the population into the sum of variance of contributing factors by using a one-way ANOVA. We define heritability as:

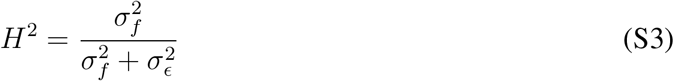

The denominator of this equation is the total phenotypic variation in the population while the numerator is the part of variation that’s due to the genetic variation in the population. Our goal is to estimate the numerator 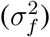 from experimental data by using a one way ANOVA.

We estimate the heritability of a quantitative trait for a collection of clonal organisms that has *N* different genotypes, and each genotype has *n_i_* members for *i* from 1 to *N* (an unbalanced clonal population). Using ANOVA, the some of squares for among genotype variation is:

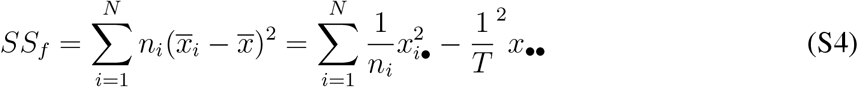

Note that *T* = Σ*n_i_* is the total number of organisms; *x_i_*_•_ is the average phenotypic value of the genotype *i* and *x*_••_ is the mean phenotype value of the whole population. Because we have *N* different genotypes, the degree of freedom is (*N* − 1). Hence the mean squares are:

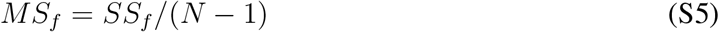

The expected value is the mean squares then is:

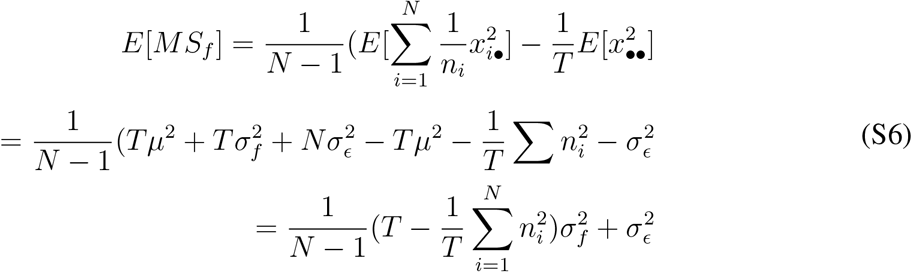

Now we calculate within-genotype variation:

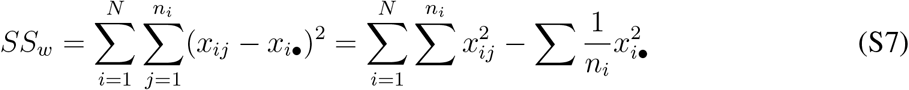

The degree of freedom here is *T* − *N* . Hence for the expected value of mean square we have:

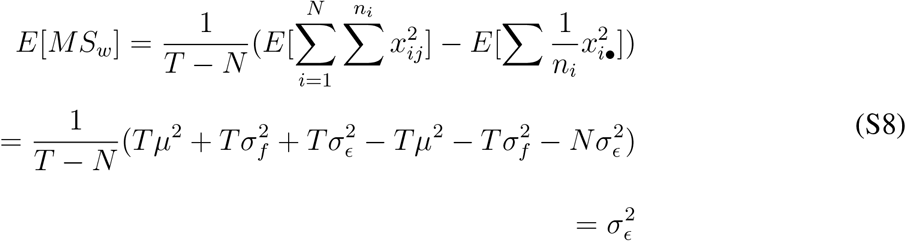

Combining the results of equations 13 and 17, we can derive an expression of 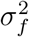:

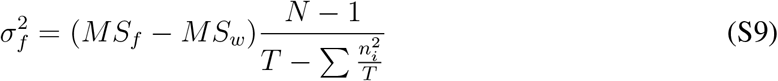

Hence the heritability is equal too:

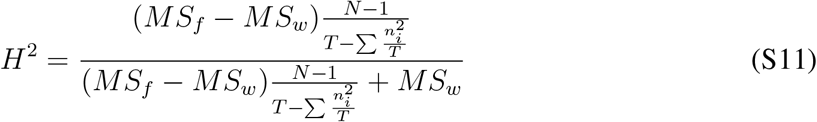

### Table of Oligonucleotides

**Table S2.**
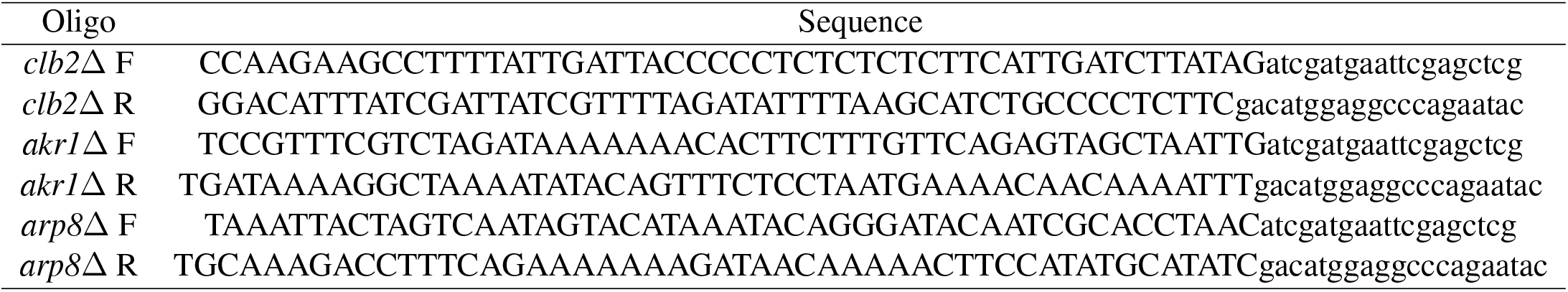
Oligonucleotides used for strain construction

**Table S2.**
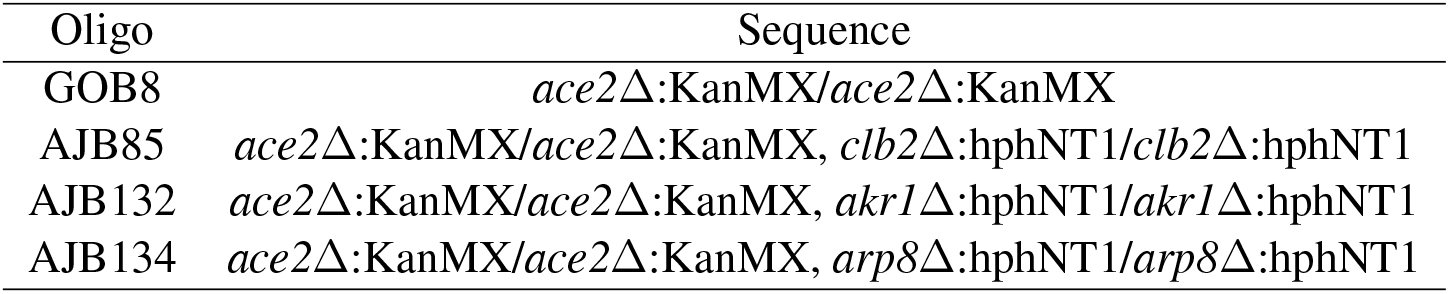
Strains

| Oligo | Sequence |
| --- | --- |
| GOB8 | *ace2*Δ:KanMX/*ace2*Δ:KanMX |
| AJB85 | *ace2*Δ:KanMX/*ace2*Δ:KanMX, *clb2*Δ:hphNT1/*clb2*Δ:hphNT1 |
| AJB132 | *ace2*Δ:KanMX/*ace2*Δ:KanMX, *akr1*Δ:hphNT1/*akr1*Δ:hphNT1 |
| AJB134 | *ace2*Δ:KanMX/*ace2*Δ:KanMX, *arp8*Δ:hphNT1/*arp8*Δ:hphNT1 |

### Supplementary Movies Supplementary Movie 1

Growth and internal stress-mediated division of a snowflake yeast cluster (genotype *ace2*∆ *clb2*∆). Sampling interval was 2.5 minutes, for a total observation time of 15 minutes.

